# Inhibitory effects of GT0918 on acute lung injury and the molecular mechanisms of anti-inflammatory response

**DOI:** 10.1101/2022.06.29.498191

**Authors:** Xiaodan Hou, Honghua Yan, Ao Wang, Cong Liu, Qianxiang Zhou, Liandong Ma, Jie Chen, Zhihua Ren, Youzhi Tong

**Author notes:** Corresponding authors: Zhihua Ren, Kintor Pharmaceutical Limited, No. 20 Songbei Road, Suzhou Industrial Park, Jiangsu, 215123, China.; Youzhi Tong, Kintor Pharmaceutical Limited, No. 20 Songbei Road, Suzhou Industrial Park, Jiangsu, 215123, China.

## Abstract

Coronavirus disease 2019 (COVID-19) has caused the public health crisis in the whole world. Anti-androgens block severe acute respiratory syndrome coronavirus 2 (SARS-CoV-2) entry and protect against severe clinical COVID-19 outcomes. GT0918, a novel androgen receptor antagonist, accelerated viral clearance and increased recovery rate in outpatients by blocking SARS-CoV-2 infection though down-regulating ACE2 and TMPRSS2 expression. Further clinical study showed that GT0918 reduced mortality rate and shortened hospital stay in hospitalized COVID-19 patients. GT0918 also exhibits protective efficacy in severe COVID-19 patient in critical care. However, the mechanism of GT0918 treatment for severe COVID-19 disease is unknown. Here, we found GT0918 decreased the expression and secretion of proinflammatory cytokines through NF-κB signaling pathway. The acute lung injury induced by LPS or Poly(I:C) was also attenuated in GT0918-treated mice, compared with vehicle control group. Moreover, GT0918 elevated the NRF2 protein level but not mRNA transcription activity. GT0918 induced proinflammatory cytokines downregulation was partially dependent on NRF2. In conclusion, our data demonstrate that GT0918 reduced cytokine release and suppressed inflammatory responses through inhibiting NF-κB signaling and activating NRF2. GT0918 is not only effective for treatment of mild to moderate COVID-19 patients, but also a potential therapeutic drug for severe COVID-19 patients by reducing the risk of cytokine storm and acute respiratory distress syndrome.

## INTRODUCTION

Coronavirus disease 2019 (COVID-19), which is caused by severe acute respiratory syndrome coronavirus 2 (SARS-CoV-2), emerged in late 2019 and became the pandemic, which threatens human health and public safety. As of December 19, 2021, over 273.3 million cases and 5.3 million deaths have been reported globally [1]. The most common symptoms of COVID-19 are fever, dry cough and tiredness. Less common symptoms include aches and pains, sore throat, diarrhea, conjunctivitis, headache, loss of taste or smell. The majority of SARS-CoV-2 infected individuals have a mild or moderate symptom and will recover in a valid time once infected. However, up to 20% of COVID-19 patients developed dyspnea one week later, and rapidly progressed to acute respiratory distress syndrome (ARDS), septic shock, multiple organ failure and even death, especially elderly patients or those who have preexisting health conditions [2]. SARS-CoV-2 recognizes angiotensin-converting enzyme 2 (ACE2) to attach to cells of the host, especially respiratory-related epithelial cells [3, 4]. SARS-CoV-2 infected cells appealed to attract immune cells into the lung, the pathogenicity involves innate immunity, T and B cell immune response and viral-neutralizing antibody response [5]. During the process, the expression of proinflammatory cytokines (TNF-α, IL-6, and IL-1β) was promoted [6]. IL-6, particularly important among the numerous increased cytokines in serum, is correlated with clinical adverse outcomes [3]. Cytokine release storm (CRS) is a common characteristic in severe COVID-19 patients. Most of the severe COVID-19 patients suffered from CRS-induced a series of side effects, such as severe lung injury and ARDS [7, 8].

According to previous studies, males are more susceptible to COVID-19 than females [9]. Compared to female, the mortality with COVID-19 was higher in males across age groups [10]. Androgen-deprivation therapies (ADTs) may protect prostate cancer patients from SARS-CoV-2 infections [11]. This predicted the role of sex hormones in antivirus and immune response. The transmembrane protease, serine 2 (TMPRSS2)-mediated cleavage of spike protein is necessary for SARS-CoV-2 virus to entry into cells [12]. A TMPRSS2 inhibitor could block the SARS-CoV-2 entry and might be an option for COVID-19 patient treatment. Androgen receptor is the only known TMPRSS2 direct regulator. Previous study showed that the antiandrogen enzalutamide downregulates TMPRSS2 expression and reduces cellular entry of SARS-CoV-2 in human lung cells [13]. Dutasteride and spironolactone, the top candidates that modulate AR signaling, were reported to reduce both ACE2 and TMPRSS2 levels in a dose-dependent manner and decreased the entry of SARS-CoV-2 [14, 15]. In males with mild COVID-19 symptoms, treatment with dutasteride reduces viral loading and inflammatory responses [16]. These studies demonstrated that anti-androgens could protect against severe clinical COVID-19 outcomes. GT0918, a second-generation androgen receptor antagonist, showed ten times more potent to antagonize AR activity than enzalutamide [17, 18]. In addition to directly inhibit AR activity, GT0918 has been shown to decrease AR expression, which is not affected by bicalutamide or enzalutamide treatment [18]. GT0918 decreased the expression of ACE2 and TMPRSS2 dose-dependently [19]. Further clinical study showed that GT0918 significantly accelerates viral clearance and reduces time to clinical remission in patients with mild to moderate COVID-19 patients versus placebo [20]. The hospitalization rate was reduced by 91% in GT0918 treated men compared to standard of care [21]. Besides, GT0918 increased recovery rate, reduced mortality rate and shortened hospital stay in hospitalized COVID-19 patients [22]. Meantime, GT0918 dramatically improved the immunologic, inflammatory, thrombotic and oxygen markers in both females and males COVID-19 outpatients. In Day 1 and Day 7 after SARS-CoV-2 infection, GT0918 reduced the numbers of neutrophils. Lymphocytes and Eosinophils were higher in the GT0918 group. The levels of CRP (C-reactive protein), D-dimer and Fibrinogen were also decreased in GT0918-treated group [23]. Since GT0918 is effective for treatment of mild to moderate COVID-19 patients. The patient number needed high flow oxygen devices, invasive mechanical ventilation, extracorporeal membrane oxygenation, vasopressors was also dramatically reduced in GT0918-treatment severe COVID-19 patients [5]. But the mechanism of GT0918 for severe COVID-19 patient treatment remain to be explored.

In the present study, we conducted lipopolysaccharide (LPS)-induced inflammation injury to explore the protective mechanism of GT0918. Compared with the control group, RNA sequencing (RNA-seq) showed the transcriptional signatures of GT0918 were enriched in immune response related signaling. GT0918 decreased the expression and secretion of proinflammatory cytokines, attenuated the acute lung injury in mice. Our data clarified the underlying mechanism of GT0918 regulation on immune response and provides a therapeutic strategy for severe COVID-19 patients.

## MATERIALS AND METHODS

### Cell culture and reagents

RAW264.7, THP-1 and U937 cell lines were purchased from the American Type Culture Collection (ATCC, United States). RAW264.7 cells were cultured in DMEM medium (HyClone, SH30022.01) with 10% FBS (BI, 04-001-1ACS), 100 U/ml penicillin and 100 μg/ml streptomycin at 37°C in a humidified 5% CO2, 95% air atmosphere. THP-1 and U937 cells were cultured in RPMI-1640 (Meilunbio, MA0215) complete medium. To acquire a macrophage-like phenotype, 200 ng/ml of PMA (Sigma, P1585) was added for 48h. Human peripheral blood mononuclear cells (hPBMCs) were purchased from ORICELLS (Shanghai, China) and cultured in RPMI-1640 complete medium. GT0918 was synthesize by Kintor Pharmaceutical limited (Suzhou, China). LPS (Sigma, L2880) or Poly(I:C) (Sigma, P9582) was used for acute lung injury induction.

### RNA-sequencing

RAW264.7 cells were pre-treated with 1μM or 3μM of GT0918 for 2h and then added 100ng/ml of LPS into the culture medium for another 16h. Cell pellets was collected and isolated for total RNA. 1 μg total RNA with RIN value above 7 was used for following library preparation. Next generation sequencing library preparations were constructed according to the manufacturer’s protocol (NEBNext® Ultra™ RNA Library Prep Kit for Illumina®). The sequences were processed and analyzed by GENEWIZ (Suzhou, China).

### RNA extraction and real-time PCR

RAW264.7 cells (or PMA-THP-1, PMA-U937, hPBMCs) were pre-treated with GT0918 for 2h and then added 100ng/ml of LPS into the culture medium for 16h. The culture medium was collected, and the cells were immediately washed with ice-cold PBS. Cell pellets was collected and isolated for total RNA with MiniBEST Universal RNA Extraction Kit (Takara, #9767). Total RNA (1 μg) was reversed to cDNA with PrimeScript™ RT reagent Kit (Takara, RR047Q), which was performed according to the manufacturer’s instructions. Real-time quantitative reactions were set up in triplicate in a 96-well plate, and each reaction contained 1 μl of cDNA and the SYBR Green PCR mix (Takara, RR420A), to which gene-specific forward and reverse PCR primers were added. The following primers were used: Human TNF-α sense: 5- CTCTTCTGCCTGCTGCACTTTG -3, antisense: 5- ATGGGCTACAGGCTTGTCACTC -3; and Human IL-6 sense: 5- AGACAGCCACTCACCTCTTCAG -3, antisense: 5- TTCTGCCAGTGCCTCTTTGCTG -3; Human IL-1β sense: 5- CCACAGACCTTCCAGGAGAATG -3, antisense: 5- GTGCAGTTCAGTGATCGTACAGG -3; human GAPDH sense: 5- GTCTCCTCTGACTTCAACAGCG -3, antisense: 5- ACCACCCTGTTGCTGTAGCCAA -3; human NRF2 sense: 5- TCTGACTCCGGCATTTCACT -3, antisense: 5- GGCACTATCTAGCTCTTCCA -3; Mouse TNF-α sense: 5- GGTGCCTATGTCTCAGCCTCTT -3, antisense: 5- GCCATAGAACTGATGAGAGGGAG -3; and Mouse IL-6 sense: 5- TACCACTTCACAAGTCGGAGGC -3, antisense: 5- CTGCAAGTGCATCATCGTTGTTC -3; Mouse IL-1β sense: 5- TGGACCTTCCAGGATGAGGACA -3, antisense: 5- GTTCATCTCGGAGCCTGTAGTG -3; Mouse GAPDH sense: 5- CATCACTGCCACCCAGAAGACTG -3, antisense: 5- ATGCCAGTGAGCTTCCCGTTCAG -3; Mouse NRF2 sense: 5- CAGCATAGAGCAGGACATGGAG -3, antisense: 5- GAACAGCGGTAGTATCAGCCAG -3; The mRNA levels of the target genes were normalized to GAPDH. The data were analyzed using GraphPad Prism 5.

### Immunoblotting

Cells were collected and total protein was extracted with RIPA lysis buffer (Beyotime, P0013B) containing 1 X protease inhibitor mixture and 1 X PMSF. The proteins were resolved on 10% SDS-polyacrylamide gels (GenScript, M00666) and transferred to polyvinylidene fluoride membranes (Merck Millipore, ISEQ00010). The membranes were blocked with 5% BSA in TBST for 1 hour. Proteins of interest were detected with specific antibodies. The antibodies used for immunoblotting (IB): p-IκBα antibody (CST, 2859s), IκBα antibody (CST, 4814s), p-p65 antibody (CST, 3033s), p65 antibody (CST, 8242s), GAPDH antibody (ABclonal, AC033), NRF2 antibody (CST, 12721T), NRF2 antibody (Abcam, ab62352). The blots were scanned using a AllianceQ9 Advance imaging system (UVETEC).

### siRNA delivery and NRF2 knockdown

Prior to electroporation, 1×10^6^ RAW264.7 cells in 100μl of EBEL buffer (Etta Biotech) were mixed with si-RNAs. Electroporation was performed at 210 V according to the manufacturer’s instructions (Etta Biotech, X-Porator H1). The sequences of siRNAs targeting mouse NRF2 were as follows: siRNA#1 sense: 5- GCAGGACAUGGAUUUGAUUTT-3, antisense: 5- AAUCAAAUCCAUGUCCUGCTT-3; siRNA#2 sense: 5- GCAGGAGAGGUAAGAAUAATT-3, antisense:5-UUAUUCUUACCUCUCCUGCTT-3.

### Acute lung injury animal model

6∼8 weeks BALB/C female mice were purchased from Beijing Vital River Laboratory Animal Technology Co., Ltd. The model of acute lung injury (ALI) was established by intranasal injection of 50µl of 4mg/ml LPS solution or 50 µL of 0.06% Poly(I:C) solution in each BALB/C female mouse after anesthesia with 2 - 5% isoflurane. Mice were intragastrically administered with GT0918 (20 mg/kg or 40 mg/kg) or the same volume of 0.5% CMC-Na solution at 16h and 1h before LPS or Poly(I:C) induction and 6h and 18h post induction. 24 h post induction, animals were sacrificed and bronchoalveolar lavage fluid (BALF) was collected. Total cell number in BALF was counted using a hemocytometer. BALF supernatant sample was transferred for TNF-α and IL-6 detection. The lung tissues were fixed for HE staining and pathologic scoring.

### ELISA

Cell supernatant or BALF was collected and diluted for detection with TNF-α ELISA kit (Multisciences, EK282-01) or IL-6 ELISA kit (Multisciences, EK206-01) according to the manufacturer’s protocol. The OD values were detected at 450nm and 570nm with Synery H1 Hybid Reader (BioTek).

### HE staining

Mice were sacrificed and the lung tissues were fixed with formalin. HE staining was conducted by Servicebio (Wuhan, China). Sections were viewed under a bright-field microscope and scored according to captured Images using a 3DHISTECH (Hungary). Pathological analysis of lung injury in mice was carried out according to the following criteria: alveolar hemorrhage, alveolar edema, infiltration of neutrophils, thickening of alveolar wall; Score based on severity of lesion was subsequently categorized as follows: 0: negative; 1: mild; 2: moderate; 3: severe; 4: extremely severe.

### Statistical analysis

The data are presented as means ± SD. Two-tailed unpaired Student’s t test was used to compare two groups, and one-way ANOVA was used for multiple comparisons. The GraphPad Prism 5 software program was used to analyze the data. All reported differences were *p < 0.05, **p < 0.01, ^#^p < 0.001 unless otherwise stated.

## RESULTS

### GT0918 reversed the LPS-induced inflammatory signature

To explore the protective mechanism of GT0918 in severe COVID-19 patient, we used LPS-induced inflammation injury model to evaluate the regulation of GT0918 on immune response. Compared with the LPS-induced group, RNA sequencing (RNA-seq) showed the transcriptional signatures of GT0918 were enriched in immune response related signaling. We pretreated RAW264.7 cells with GT0918, then add LPS to the mediums for overnight and conducted RNA sequencing. Transcriptome analysis was performed to compare the genes expression with or without GT0918 treatment. The LPS induction changed the gene expression profile markedly, GT0918 reversed most of the differentially expressed genes (DEGs) induced by LPS (Figure 1A). While the levels of 299 genes were notably downregulated, the expressions of 60 genes were upregulated more than 2-folds in 3μM of GT0918 group (Figure 1B). KEGG (Kyoto Encyclopedia of Genes and Genomes) enrichment showed the DEGs were related to inflammatory-related disease, especially in virus infection, some of them were also correlated with COVID-19 (Figure 1C). Immune-related signaling such as TNF signaling pathway, JAK-STAT signaling pathway and NF-κB signaling pathway were also enriched in GT0918 treatment group (Figure 1D). KEGG facilitates functional pathway enrichment to determine the involved physiological activities of these DEGs. A variety of immune-related pathway and inflammatory cytokines were involved (Figure 1E). After GT0918 treatment, most of the proinflammatory cytokines were reduced dose-dependently (Figure 1F). Continuous release of pro-inflammatory factors might lead to CRS, a common syndrome in severe COVID-19 patients. GT0918 induced down-regulation of pro-inflammatory cytokines expression may keep COVID-19 patients avoid of CRS.

**Figure 1.**
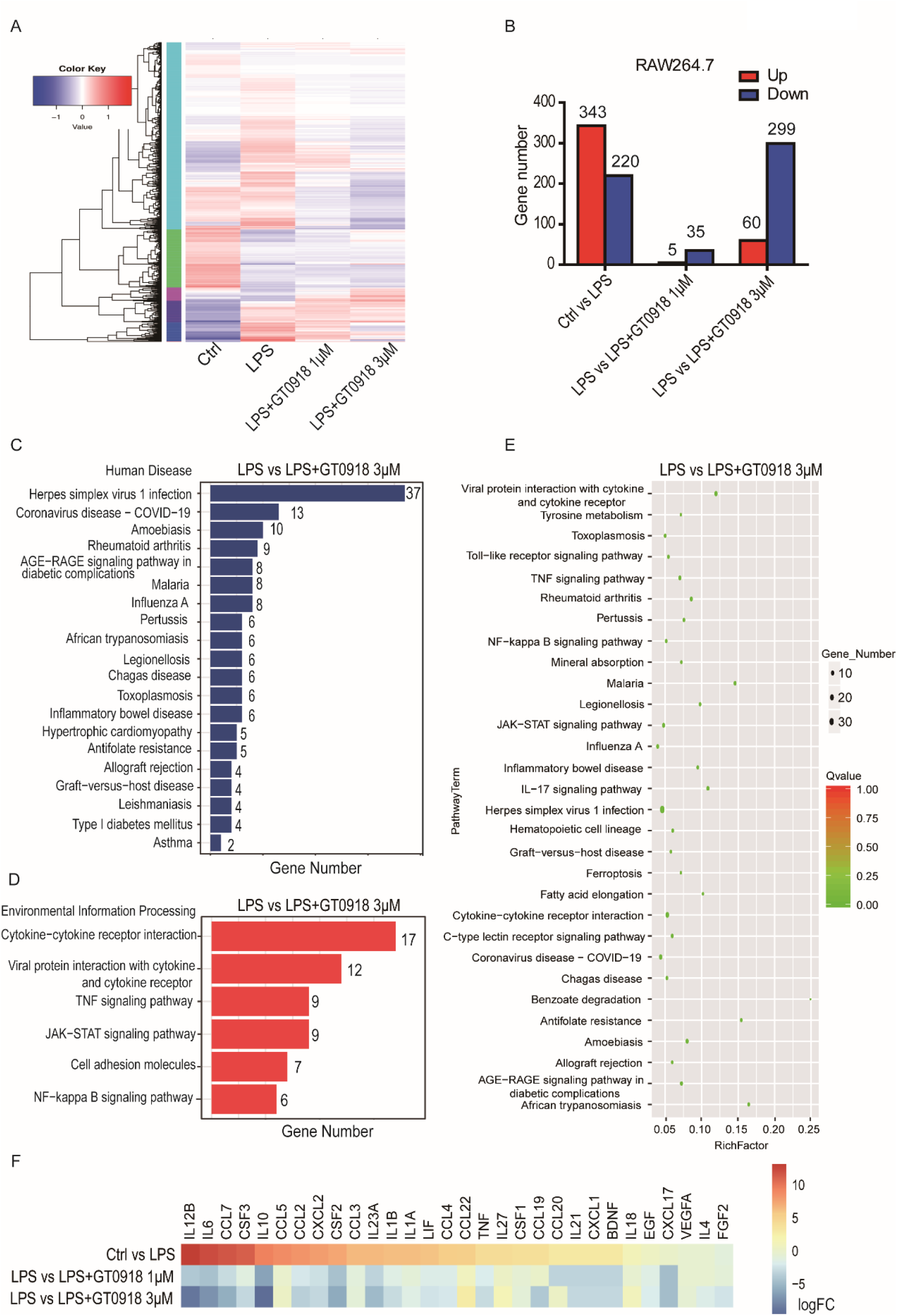
LPS-induced inflammatory signature was reversed by GT0918. **A**. Transcriptome sequencing of LPS-induced model pretreated with or without GT0918 in RAW264.7 cells. Gene expression profiles were performed with log10 (RPKM+1) value, with high expression gene in red and low expression gene in blue. Color from blue to red indicates higher gene expression. **B**. Summary of gene numbers that are significantly upregulated or downregulated between groups. **C**. Kyoto encyclopedia of genes and genomes (KEGG) enrichment of DEGs in human disease between LPS and LPS+GT0918-3μM group. **D**. KEGG analysis of DEGs involved in environmental information processing between LPS and LPS+GT0918-3μM group. **E**. KEGG analysis facilitates functional pathway enrichment of DEGs. The size of the bubble means gene number; the color depth means Q-value, and the rich factor means the gene number/the total gene number in the y-axis item. **F**. Immune response related cytokines and chemokines with significant changes between groups. Upregulated gene in red and downregulated gene in blue, color from blue to red indicates higher fold change.

### GT0918 decreased the proinflammatory cytokines expression through NF-κ B signaling

To verify the effect of GT0918 on the expression of proinflammatory cytokines, we used monocyte/macrophage cell lines and human PBMCs (Peripheral Blood Mononuclear Cells) to detect the changes of LPS-induced IL-6, TNF-α and IL-1β under GT0918 treatment. In RAW264.7 and U937 cells, GT0918 reduced the expression of IL-6, TNF-α and IL-1β dose-dependently (Figure 2A). In THP-1 cells, GT0918 decreased the expression of IL-6 and TNF-α, IL-1β was slightly increased. In human PBMCs, GT0918 reduced the expression of IL-6, TNF-α dose-dependently. IL-1 β was mild upregulated followed by significant inhibition with different concentration of GT0918 treatment. The secretion of IL-6 and TNF-α was also inhibited dose-dependently (Figure 2B). As we know, NF-κB signaling induces several pro-inflammatory factors. We explored whether the downregulation of these proinflammatory cytokines was through NF-κB signaling pathway. The data showed GT0918 downregulated the activation of p65 by decreasing phosphorylation of IκBα, and then inhibited the activation of NF-κB pathway in a dose-dependent manner, suggesting GT0918 decreased the proinflammatory cytokines expression through NF-κB signaling (Figure 2C).

**Figure 2.**
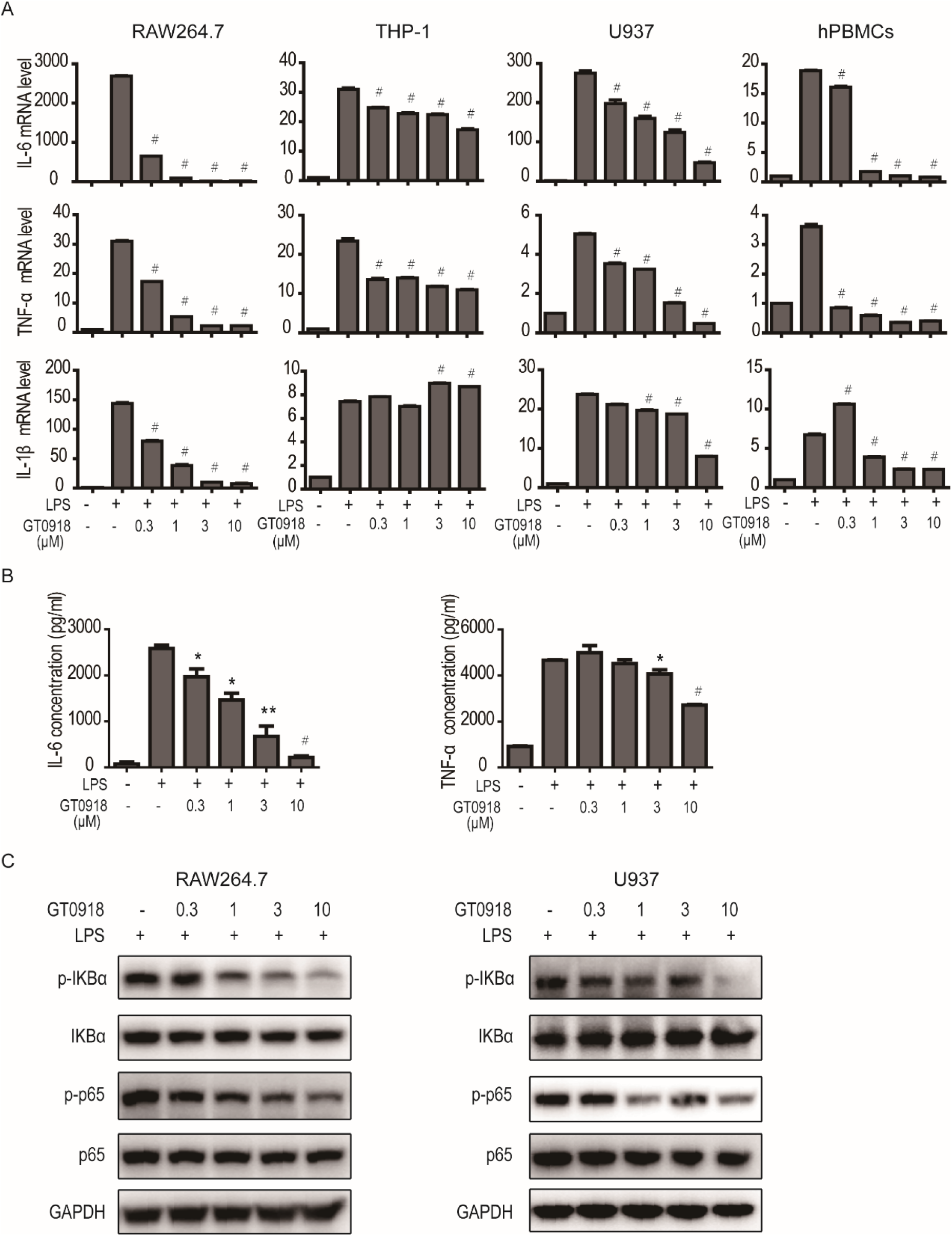
GT0918 decreased the expression and release of proinflammatory cytokines and attenuated NF-κB signaling dose-dependently. **A**. The expression of IL-6, TNF-α and IL-1β under different concentration of GT0918 treatment in RAW264.7, THP-1, U937 cells and hPBMCs. RAW264.7 cells (or PMA-THP-1, PMA-U937, hPBMCs) were pre-treated with GT0918 for 2h and then added 100ng/ml of LPS into the culture medium for 16h. **B**. The concentration of IL-6 and TNF-α in the supernatant of RAW264.7 cells under GT0918 treatment. **C**. The effect of GT0918 on NF-κB signaling in RAW264.7 and U937 cells. RAW264.7 or PMA-U937 cells were pre-treated with GT0918 for 2h and then added 100ng/ml of LPS into the culture medium for 0.5h.

### GT0918 attenuated LPS-induced acute lung injury in vivo

We conducted in vivo studies to determine the effect of GT0918 on LPS-induced acute lung injury. Our results demonstrated that LPS stimulation increased the total cell number of BALF (Bronchoalveolar Lavage Fluid). The cell type of BALF in LPS-induced mouse model includes lymphocyte, neutrophil, monocyte/ macrophage and a small number of other cell types. After GT0918 treatment, the total cell number of BALF was lower more than three times than LPS group (Figure 3A). In a subset of immune cells, lymphocyte was the most obviously declined (Figure 3B). Neutrophil and monocyte/ macrophage were also reduced with GT0918 treatment, but there is no statistical difference because of the variation in LPS group (Figure 3C&3D). ELISA analysis of BALF showed the concentration of TNF-α and IL-6 were both reduced with GT0918 treatment (Figure 3E&3F). This result was consistent with GT0918 anti-inflammatory response *in vitro*. HE staining demonstrated LPS induced the irregular alveolar structure, more neutrophil infiltration and thickened alveolar septa of lung injury in mice. GT0918 attenuated LPS-induced acute lung injury dose-dependently (Figure 3G&3H).

**Figure 3.**
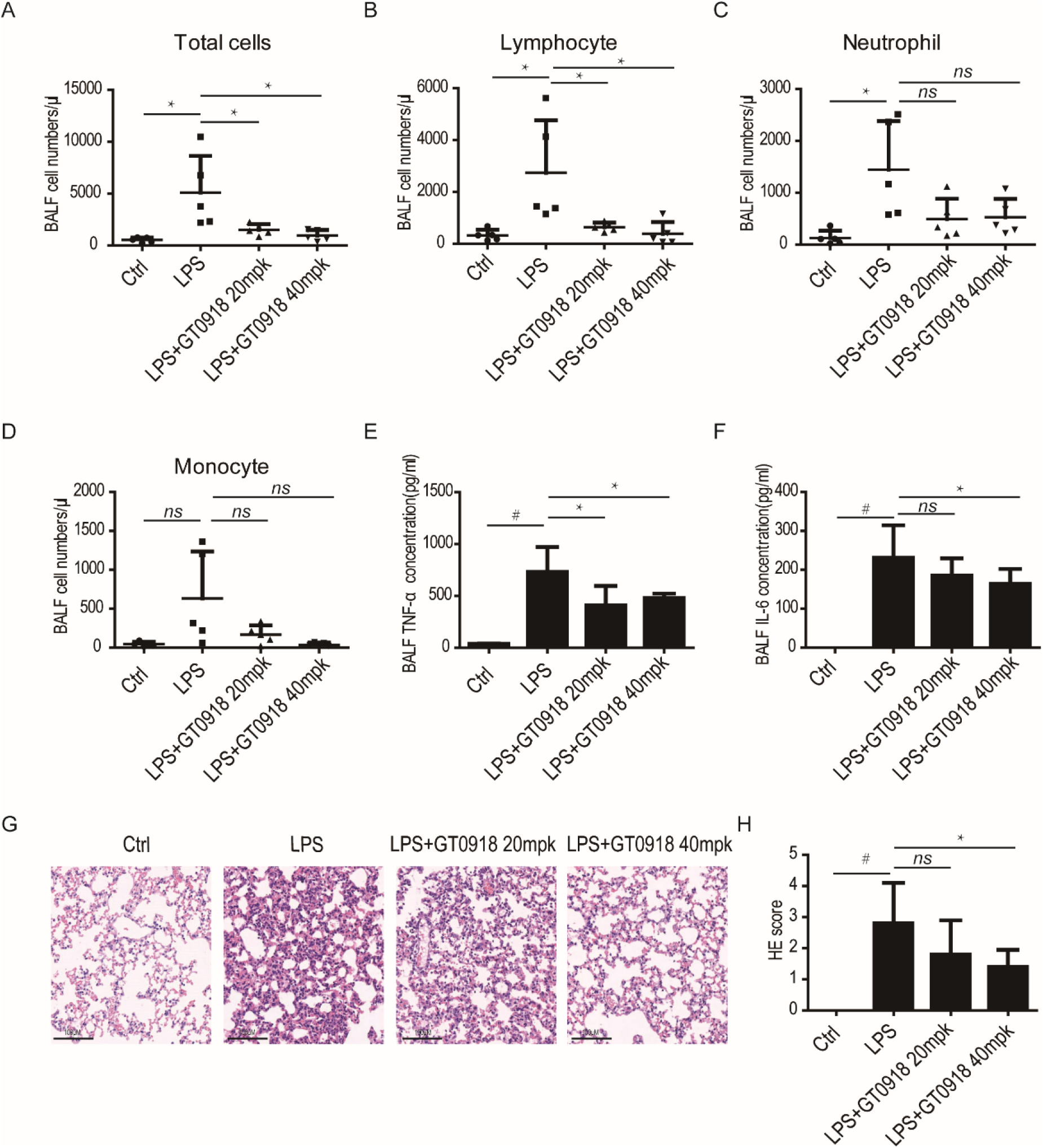
GT0918 attenuated LPS-induced acute lung injury *in vivo*. **A-D**. The effect of GT0918 on the immune cell numbers in BALF of LPS-induced mouse model. Mice were intragastrically administered with GT0918 (20 mg/kg or 40 mg/kg) or the same volume of 0.5% CMC-Na solution at 16h and 1h before LPS induction and 6h and 18h post induction. **E**. The concentration of TNF-α in the BALF of LPS-induced mouse model. **F**. The concentration of IL-6 in the BALF of LPS-induced mouse model. **G**. HE staining showed the effect of GT0918 on the LPS-induced lung injury. **H**. HE scores based on severity of lesion was analyzed as follows: 0: negative; 1: mild; 2: moderate; 3: severe; 4: very severe.

### GT0918 attenuated Poly(I:C)-induced acute lung injury

Since SARS-CoV-2 is RNA virus, we tried to conducted Poly(I:C) -induced model to mimic the acute lung injury of SARS-CoV-2 infection *in vivo*, which is a mismatched double-stranded RNA. Poly(I:C) stimulation increased the immune cell number of BALF, mainly neutrophil and monocyte, only a few lymphocytes were involved (Figure 4A). There was a trend of monocyte decrease with GT0918 treatment, but there is no statistical difference because of the variation in Poly(I:C) group (Figure 4B). GT0918 extremely declined the number of neutrophils in BALF (Figure 4C). The concentration of TNF-α and IL-6 were both decreased with GT0918 treatment (Figure 4D&4E). HE staining showed Poly(I:C)induced more neutrophil infiltration in the lung tissue of mice. 40mpk of GT0918 attenuated neutrophil infiltration. The thickened alveolar septa induced by Poly(I:C) was also attenuated with GT0918 treatment (Figure 4F&4G). The results demonstrated that GT0918 attenuated both LPS and Poly(I:C)-induced acute lung injury *in vivo*.

**Figure 4.**
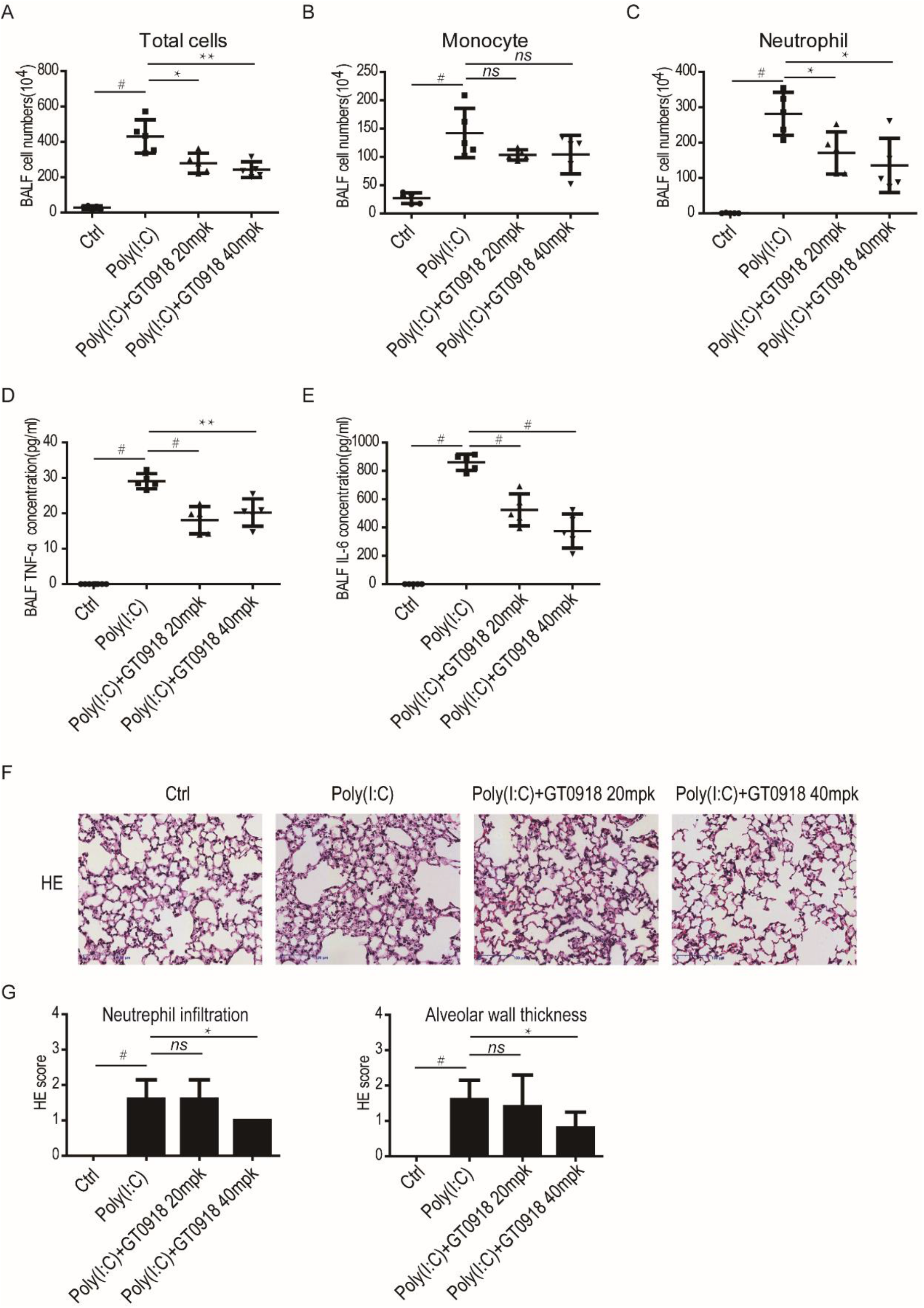
GT0918 attenuated Poly(I:C)-induced acute lung injury *in vivo*. **A-C**. The effect of GT0918 on the immune cell numbers in BALF of Poly(I:C)-induced mouse model. Mice were intragastrically administered with GT0918 (20 mg/kg or 40 mg/kg) or the same volume of 0.5% CMC-Na solution at 16h and 1h before Poly(I:C) induction and 6h and 18h post induction. **D**. The concentration of TNF-α in the BALF of Poly(I:C)-induced mouse model. **E**. The concentration of IL-6 in the BALF of Poly(I:C)-induced mouse model. **F**. HE staining showed the effect of GT0918 on the Poly(I:C)-induced lung injury. **G**. HE scores of neutrophil infiltration and alveolar septa thickness was analyzed as follows: 0: negative; 1: mild; 2: moderate; 3: severe; 4: very severe.

### GT0918 activated NRF2 signaling

By screening of more than 200 transcription factors, we found the DNA-binding activity of NRF2 (NFE2L2) was enhanced with GT0918 treatment (data not shown). NRF2 regulates antioxidant defense signaling and negatively regulates NF-κB signaling [24]. Further analysis showed the protein level of NRF2 was gradually increased with different dose of GT0918 treatment with or without LPS stimulation (Figure 5A&5B). But the mRNA level of NRF2 was not changed within 3h or 18h treatment of GT0918 in RAW264.7. At 10μM of GT0918, NRF2 mRNA level was slightly downregulated (Figure 5C). In U937 cells, NRF2 mRNA level was also not changed with different concentration of GT0918 treatment (Figure 5D). With NRF2 protein level increased, its target protein level of HO-1 was also increased (Figure 5E). These results showed that GT0918 was not only an androgen receptor antagonist, but also participated in NRF2 protein regulation.

**Figure 5.**
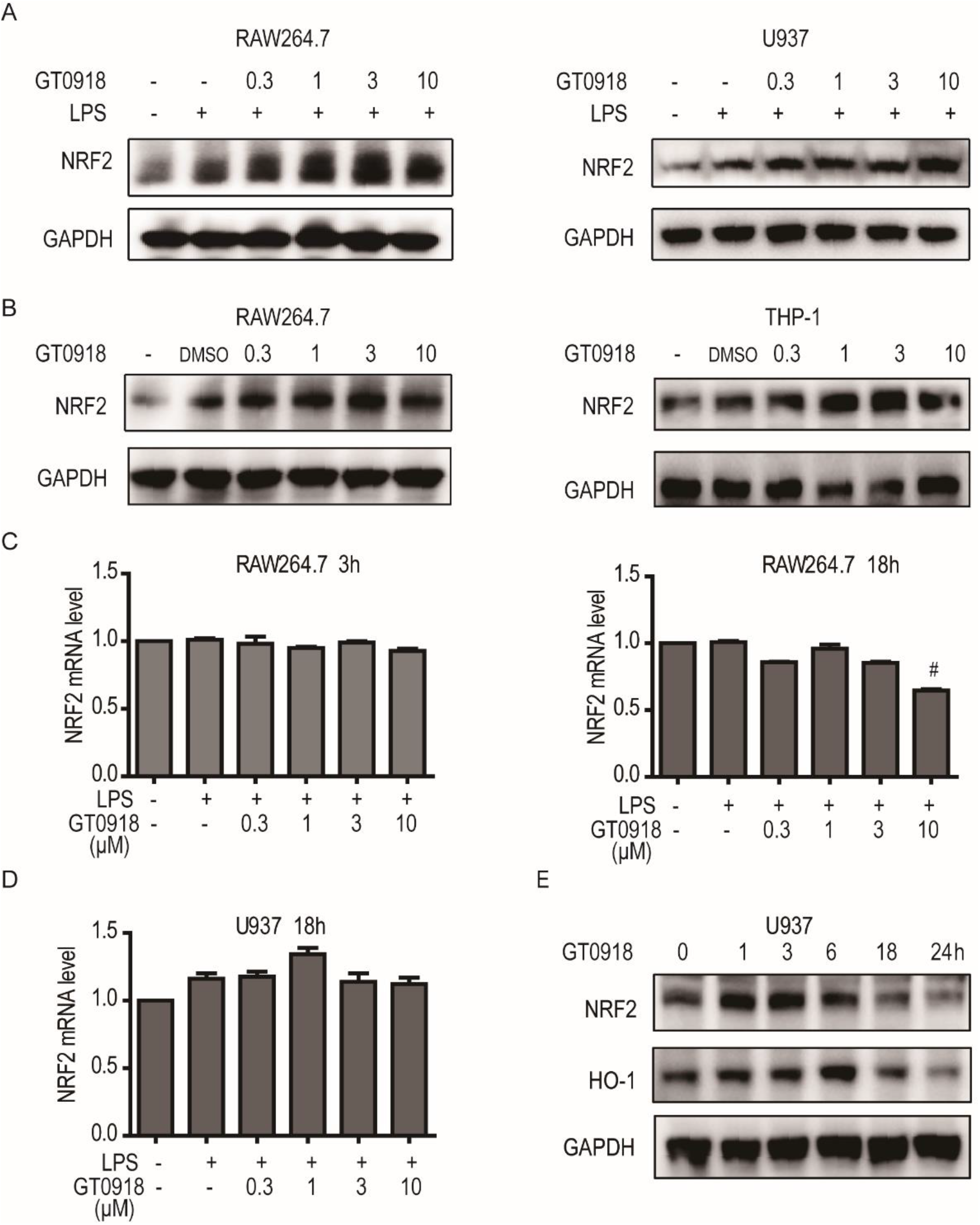
GT0918 activated NRF2 signaling. A. The protein level of NRF2 with different dose of GT0918 treatment under LPS stimulation. RAW264.7 or PMA-U937 cells were pre-treated with GT0918 for 2h and then added 100ng/ml of LPS into the culture medium for 1h. B. The protein level of NRF2 with different dose of GT0918 treatment. RAW264.7 or PMA-THP-1 cells were treated with GT0918 for 3h. C. The mRNA level of NRF2 with GT0918 treatment under LPS stimulation in RAW264.7 cells. RAW264.7 cells were pre-treated with different dose GT0918 for 2h and then added 100ng/ml of LPS into the culture medium for 1h or 16h. D. The mRNA level of NRF2 with GT0918 treatment in U937 cells. PMA-U937 cells were pre-treated with GT0918 for 2h and then added 100ng/ml of LPS into the culture medium for 16h. E. The protein level of NRF2 and HO-1 with GT0918 treatment in U937 cells. PMA-U937 cells were treated with 10μM of GT0918 for the indicated time.

### GT0918 induced proinflammatory cytokines downregulation was dependent on NRF2

Next, we synthesize siRNAs to specifically knockdown NRF2 and explore whether GT0918 induced proinflammatory cytokines downregulation was dependent on NRF2. By electroporation of siRNAs into RAW264.7 cells, NRF2 mRNA and protein levels were both downregulated (Figure 6A). The mRNA levels of IL-6, TNF-α and IL-1β were decreased with GT0918 treatment. Moreover, GT0918-induced IL-6, TNF-α and IL-1β downregulation was reversed in NRF2 knockdown RAW264.7 cells (Figure 6B&6C&6D). Compared to control, the concentration of TNF-α and IL-6 were also reversed in NRF2 knockdown RAW264.7 cells under GT0918 treatment (Figure 6E&6F). GT0918 decreased the proinflammatory cytokines levels through NF-κB signaling. In NRF2 knockdown RAW264.7 cells, the downregulation of p65 phosphorylation was reversed with GT0918 treatment. These results demonstrated that GT0918 induced proinflammatory cytokines downregulation was dependent on NRF2.

**Figure 6.**
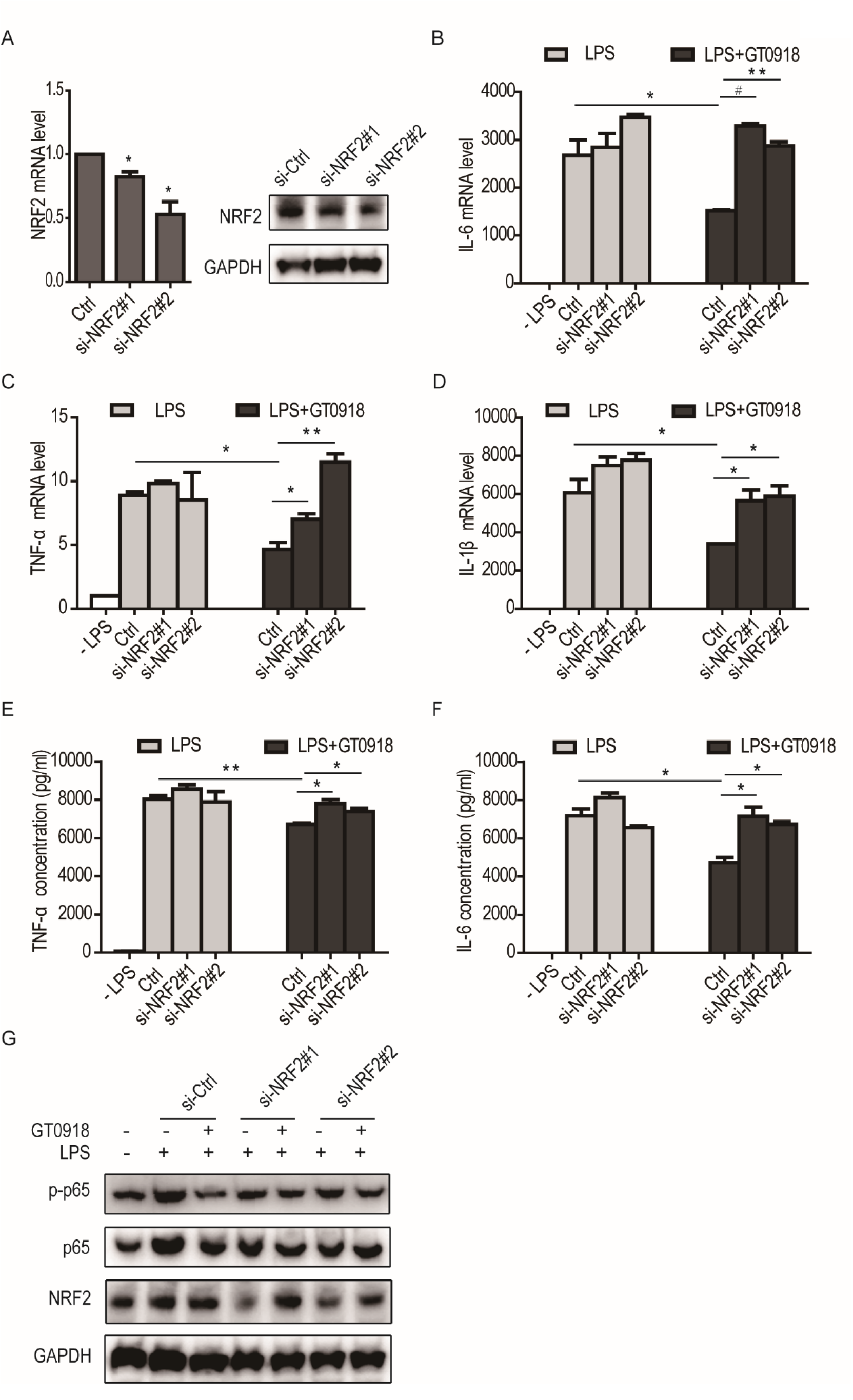
GT0918 induced proinflammatory cytokines downregulation was dependent on NRF2. **A**. The mRNA and protein levels of NRF2 in control or NRF2 knockdown RAW264.7 cells. **B-D**. The mRNA levels of IL-6 (**B**), TNF-α (**C**) and IL-1β (**D**) in control or NRF2 knockdown RAW264.7 cells. After electrotransfected with control or NRF2 si-RNAs for 24h, RAW264.7 cells were pre-treated with 3μM of GT0918 for 2h and then added 100ng/ml of LPS into the culture medium for 16h. **E&F**. The concentration of TNF-α (**E**)and IL-6 (**F**)in control or NRF2 knockdown RAW264.7 cell supernatants. **G**. The effect of GT0918 on NF-κB signaling in control or NRF2 knockdown RAW264.7 cells. After electrotransfected with control or NRF2 si-RNAs for 24h, RAW264.7 cells were pre-treated with 3μM of GT0918 for 2h and then added 100ng/ml of LPS into the culture medium for 0.5h.

## DISCUSSION

COVID-19 is rapidly spreading all around the world. Global scientists have made great progress in developing medicines and vaccines against COVID-19. Vaccines and neutralizing antibodies have become powerful tools to control the epidemic, leading to a slow increase in the number of SARS-CoV-2 infections. Unexpected emerging of SARS-CoV-2 virus variants, which carrying multiple mutations on spike protein binding domain, may be resistant to the existing treatments or vaccines. Oral drugs are more important than monoclonal antibodies or vaccines for long-term control of outbreaks. Compared with the monoclonal antibodies, oral drugs are easy to produce, and convenient to use. Up to now, only Merck’s Molnupiravir and Pfizer’s Paxlovid have been approved to treat people with mild-to-moderate COVID-19. Both drugs are antiviral medications through inhibiting SARS-CoV-2 replication. ADT is associated with decreased SARS-CoV-2 infection and severity of COVID-19 in men [25]. Previous reports showed GT0918, a novel androgen receptor antagonist, was effective for accelerated viral clearance, increased recovery rate and reduced mortality rate of COVID-19 patients [20-23]. In this study, we found GT0918 decreased the expression of TNF-α, IL-6, and IL-1β. These proinflammatory factors are quickly eliminated in mild-to-moderate patients, but their delayed and continued elevation can lead to cytokine release storm (CRS). CRS induced a series of side effects, such as severe lung injury and ARDS, is a common characteristic in severe COVID-19 patients. GT0918 decreased the levels of these proinflammatory cytokines dose-dependently, protected against CRS-induced severe clinical COVID-19 outcomes. Further analysis indicated GT0918 downregulated the activation of p65 by decreasing phosphorylation of IκBα, and then inhibited the activation of NF-κB pathway in a dose-dependent manner, suggesting GT0918 inhibited proinflammatory cytokines expression through NF-κB signaling. LPS or Poly(I:C) induced acute lung injury was conducted to mimic the effect of SARS-CoV-2 on the lung tissue *in vivo*. We found the immune cells in BALF were declined in GT0918 treatment mice. The release of TNF-α and IL-6 was also significantly decreased at 40mpk of GT0918. Inflammatory infiltration in injured lung tissue was reduced and the destruction of alveolar structures was mild.

Moreover, by screening of more than 200 transcription factors, we found the DNA-binding activity of NRF2 was enhanced with GT0918 treatment. GT0918 is not only an androgen receptor antagonist and degrader, but also participated in the regulation of NRF2. We found the protein level of NRF2 was gradually increased, but the mRNA transcription levels were not changed within short or longer time treatment with GT0918. The results showed that GT0918 may prevent NRF2 degradation and improved NRF2 stability. In basal conditions, KEAP1 (Kelch ECH associating protein 1) binds to NRF2 and promotes its degradation by the canonical ubiquitin-proteasome pathway [26]. The accumulated NRF2 translocated into the nucleus and initiated the target gene transcription via the antioxidant response element (ARE), such as HO-1. NRF2 antioxidant gene expression pathway is suppressed in COVID-19 patients. NRF2 agonists 4-OI and DMF is broadly effective in limiting SARS-CoV-2 virus replication and suppressing the proinflammatory responses [27]. GT0918 activated NRF2 mechanism predicted the involvement of the NRF2 antioxidant pathway in the protection of SARS-CoV-2-induced immune response. When NRF2 was knocked down, GT0918 induced proinflammatory cytokines downregulation was partially reversed. This demonstrated that GT0918 regulated immune response was through NF-κB pathway and NRF2 antioxidant pathway. Our data revealed the underlying mechanism of GT0918 regulation on immune response. Thus, GT0918 is not only effective for treatment of patients with mild to moderate COVID-19 disease, but also a potential therapeutic drug for severe COVID-19 disease.

## ACKNOWLEDGEMENT

This study was supported by National Key Research and Development Program of China (2021YFC0864700).

## REFERENCES

1. World Health Organization, Coronavirus disease 2019 (COVID-19) Situation Report – December 19, 2021;. Available from: https://www.who.int/emergencies/diseases/novel-coronavirus-2019/situation-reports.

2. Moore, J.B. and C.H. June, Cytokine release syndrome in severe COVID-19. Science, 2020. 368(6490): p. 473–474.

3. Letko, M., A. Marzi, and V. Munster, Functional assessment of cell entry and receptor usage for SARS-CoV-2 and other lineage B betacoronaviruses. Nat Microbiol, 2020. 5(4): p. 562–569.

4. Huang, Y., et al., Structural and functional properties of SARS-CoV-2 spike protein: potential antivirus drug development for COVID-19. Acta Pharmacol Sin, 2020. 41(9): p. 1141–1149.

5. Blanco-Melo, D., et al., Imbalanced Host Response to SARS-CoV-2 Drives Development of COVID-19. Cell, 2020. 181(5): p. 1036–1045 e9.

6. La Torre, F., et al., Immunological basis of virus-host interaction in COVID-19. Pediatr Allergy Immunol, 2020. 31 Suppl 26: p. 75–78.

7. Darif, D., et al., The pro-inflammatory cytokines in COVID-19 pathogenesis: What goes wrong? Microb Pathog, 2021. 153: p. 104799.

8. Tay, M.Z., et al., The trinity of COVID-19: immunity, inflammation and intervention. Nat Rev Immunol, 2020. 20(6): p. 363–374.

9. Bwire, G.M., Coronavirus: Why Men are More Vulnerable to Covid-19 Than Women? SN Compr Clin Med, 2020: p. 1–3.

10. Nguyen, N.T., et al., Male gender is a predictor of higher mortality in hospitalized adults with COVID-19. PLoS One, 2021. 16(7): p. e0254066.

11. Brookman-May, S.D. and M. May, Re: Androgen-deprivation Therapies for Prostate Cancer and Risk of Infection by SARS-CoV-2: A Population-based Study (n=4532). Eur Urol, 2020. 78(6): p. 930–931.

12. Hoffmann, M., et al., SARS-CoV-2 Cell Entry Depends on ACE2 and TMPRSS2 and Is Blocked by a Clinically Proven Protease Inhibitor. Cell, 2020. 181(2): p. 271–280 e8.

13. Leach, D.A., et al., The antiandrogen enzalutamide downregulates TMPRSS2 and reduces cellular entry of SARS-CoV-2 in human lung cells. Nat Commun, 2021. 12(1): p. 4068.

14. Goren, A., et al., Anti-androgens may protect against severe COVID-19 outcomes: results from a prospective cohort study of 77 hospitalized men. J Eur Acad Dermatol Venereol, 2021. 35(1): p.e13–e15.

15. Ghazizadeh, Z., et al., Androgen Regulates SARS-CoV-2 Receptor Levels and Is Associated with Severe COVID-19 Symptoms in Men. bioRxiv, 2020.

16. Cadegiani, F.A., et al., Early Antiandrogen Therapy With Dutasteride Reduces Viral Shedding, Inflammatory Responses, and Time-to-Remission in Males With COVID-19: A Randomized, Double-Blind, Placebo-Controlled Interventional Trial (EAT-DUTA AndroCoV Trial - Biochemical). Cureus, 2021. 13(2): p. e13047.

17. Qu, F., et al., Metabolomic profiling to evaluate the efficacy of proxalutamide, a novel androgen receptor antagonist, in prostate cancer cells. Invest New Drugs, 2020. 38(5): p. 1292–1302.

18. Zhou, T., et al., Preclinical profile and phase I clinical trial of a novel androgen receptor antagonist GT0918 in castration-resistant prostate cancer. Eur J Cancer, 2020. 134: p. 29–40.

19. Siqi Wu1*, et al., Suppression of Androgen Receptor (AR)-ACE2/TMPRSS2 Axis by AR Antagonists May Be Therapeutically Beneficial for Male COVID-2019 Patients.

20. Cadegiani, F.A., et al., Proxalutamide Significantly Accelerates Viral Clearance and Reduces Time to Clinical Remission in Patients with Mild to Moderate COVID-19: Results from a Randomized, Double-Blinded, Placebo-Controlled Trial. Cureus, 2021. 13(2): p. e13492.

21. McCoy, J., et al., Proxalutamide Reduces the Rate of Hospitalization for COVID-19 Male Outpatients: A Randomized Double-Blinded Placebo-Controlled Trial. Front Med (Lausanne), 2021. 8: p. 668698.

22. Cadegiani, F.A., et al., Final Results of a Randomized, Placebo-Controlled, Two-Arm, Parallel Clinical Trial of Proxalutamide for Hospitalized COVID-19 Patients: A Multiregional, Joint Analysis of the Proxa-Rescue AndroCoV Trial. Cureus, 2021. 13(12): p. e20691.

23. Flávio Adsuara Cadegiani, et al., Proxalutamide Improves Inflammatory, Immunologic, and Thrombogenic Markers in Mild-to-Moderate COVID-19 Males and Females: an Exploratory Analysis of a Randomized, double-Blinded, Placebo-Controlled Trial Early Antiandrogen Therapy (EAT) with Proxalutamide (The EAT-Proxa Biochemical AndroCoV-Trial). medRxiv preprint, 2021.

24. Wardyn, J.D., A.H. Ponsford, and C.M. Sanderson, Dissecting molecular cross-talk between Nrf2 and NF-kappaB response pathways. Biochem Soc Trans, 2015. 43(4): p. 621–6.

25. Kyung Min Lee, et al., A population-level analysis of the protective effects of androgen deprivation therapy against COVID-19 disease incidence and severity. Frontiers in Medicine, 2022.

26. Kansanen, E., et al., The Keap1-Nrf2 pathway: Mechanisms of activation and dysregulation in cancer. Redox Biol, 2013. 1: p. 45–9.

27. Olagnier, D., et al., SARS-CoV2-mediated suppression of NRF2-signaling reveals potent antiviral and anti-inflammatory activity of 4-octyl-itaconate and dimethyl fumarate. Nat Commun, 2020. 11(1): p. 4938.

